# Microglia Piezo1 aggravates cerebral ischemia and reperfusion injury by chemotactic recruitment of T lymphocytes

**DOI:** 10.64898/2026.01.12.699156

**Authors:** Hui-Nan Zhang, Hui-Min Chang, Bao Wang, Tian Gao, Hui-Feng Zhang, Si-Jia Xu, Xing Wang, Fei Liu, Jing Huang

## Abstract

Cerebral ischemia reperfusion injury (CIRI) is a complex pathophysiological process involving multiple mechanisms. Piezo1, a stretch-activated ion channel, has recently emerged as a potential regulator of cellular responses to ischemic conditions. However, its role in specific cells during ischemic events is not well elucidated. Here, we showed that after experimentally induced CIRI, Piezo1 channel was highly expressed in peri-infarct area, where is upregulated and activated mainly in microglia. Behavioral tests, and infarct volume measurements demonstrated that conditional Piezo1 deficient in microglia markedly ameliorated the neurological deficits and reduced the infarct volumes. Flow cytometry and immunofluorescence staining showed decreased peripheral immune cells, especially T lymphocytes infiltration into the brain in microglia Piezo1 deficient mice after stroke. Multiplex chemokine immunoassay and T cell migration assays screened that Piezo1 deficient microglia blocked the CXC chemokine ligand 10 (CXCL10) release from astrocyte, which lead to the inhibited T cell recruitment in transwell co-cultured system. Furthermore, RNA-Sequencing analysis showed that defective Piezo1 in microglia lead to its polarization towards an anti-inflammatory phenotype in response to cerebral ischemia and reduced the secretion of tumor necrosis factor-α (TNF-α) and interferon-γ (IFN-γ). Collectively, Piezo1 deficiency inhibited the secretion of TNF-α and IFN-γ in microglia. The reduction of these inflammatory cytokines further blocked the CXCL10 release from astrocyte, and ultimately ameliorated the infiltration of peripheral T lymphocytes after stroke. Our results therefore highlight the critical role of Piezo1 in microglia-orchestrated neuroinflammation and suggest a potential means for reducing stroke-induced neurological injury.

## 1. Introduction

Ischemic stroke constitutes over 70% of all strokes and is one of the leading causes of morbidity and mortality worldwide [1]. Timely restoration of cerebral blood flow is critical for effective treatment, however, this process can paradoxically aggravate brain injury, a phenomenon known as cerebral ischemia and reperfusion injury (CIRI) [2]. Peripheral leukocytes invade into the brain following ischemic stroke and play a critical role in exacerbating brain edema and further brain damage [3]. The crosstalk between the central nerves system (CNS) and peripheral immune system is regulated by a complex network. Targeting this interaction offers a promising strategy for developing effective therapies to mitigate brain injury after CIRI [4].

Microglia are inherent immune cells in CNS and play crucial roles under pathological conditions such as ischemic stroke [5, 6]. Microglia are activated and act as the earliest cells responders to CIRI [7]. Activated microglia induce the leukocytes infiltration by secreting inflammatory cytokines such as tumor necrosis factor-α (TNF-α) and interferon-γ (IFN-γ) [7]. Microglia express several mechanosensory ion channels, which involved in sensing the intense mechanical changes, thereby regulating the intracellular Ca^2+^ homeostasis [8, 9]. Mechanosensory ion channels are reported to be activated and identified as a potential trigger for inflammatory in microglia during CIRI, microglia undergo swelling, leading to changes in membrane tension [10]. However, the role of mechanosensory ion channels following stroke is still not fully clarified and warrants further research.

Piezo1 is an evolutionarily conserved mechanosensory ion channel protein [11]. Several researches have shown that Piezo1 play a critical role in pathological processes of the brain diseases [12–14]. In a mouse model of Alzheimer’s disease, Piezo1 is upregulated and activated in microglia, leading to influx of extracellular Ca^2+^ and stimulated Ca^2+^ signaling [15]. Therapeutic strategies targeting Piezo1 have been demonstrated effective in models of Alzheimer’s disease, multiple sclerosis, and glaucoma [15–17]. Specifically, microglial Piezo1 appears to play a central role in initiating neuroinflammatory responses across diverse neuropathologies [18]. The molecular mechanisms underlying Piezo1 activation triggers the Ca^2+^ influx, which subsequently stimulates downstream signaling pathways, including Ca^2+^/ calmodulin dependent protein kinase, which play a key role in the regulation of neuroinflammation [19]. Clinically, a correlation study has revealed a significant association between Pizeo1 expression in red blood cells and ischemic stroke outcome in patients [20]. Moreover, Piezo1 is observed to be upregulated in rodent stroke model [21], which have linked Piezo1 protein to cerebral ischemia. Despite these findings, the specific role of Piezo1 in mediating brain injury following CIRI remains largely unexplored and requires further investigation.

In this study, we found that Piezo1 was upregulated and activated in microglia in a mouse model of CIRI. We demonstrated that microglial Piezo1 deficiency significantly reduces neuroinflammatory responses induced by ischemic brain injury. Our results further revealed that Piezo1 activation in microglia promotes the secretion of pro-inflammatory cytokines, such as TNF-α and IFN-γ, which in turn stimulate CXC chemokine ligand 10 (CXCL10) production. This cascade exacerbates T cell infiltration and worsens brain injury. Thus, our study suggests that targeting microglial Piezo1 may offer a potential therapeutic approach for mitigating CIRI.

## 2. Methods

### 2.1. Animals

Male mice, aged 8-12 weeks old were used for this research. Mouse strains were generated and maintained on the C57BL/6 background. Microglia *Piezo1* conditional knockout (Piezo1^CKO^) mice were generated by mating *Piezo1*^flox/flox^ (Cyagen Incorporated, Suzhou, China) and CX3C chemokine receptor 1 (Cx3cr1)^CreER^ (Cyagen Incorporated) mice. The mice were intraperitoneally injected with tamoxifen (75 mg/kg, Sigma-Aldrich, St Louis, MO, USA) dissolved in corn oil intraperitoneally for 5 consecutive days 14 days before MCAO/R surgery. *Piezo1*^flox/flox^ mice were used as the control group. Behavior deficits and infarct volume were designated as the primary endpoints of the study. The number of mice used in each experiment is specified in the corresponding figure bar. All mice were housed under 12 h light/dark cycle house at 25 °C and had free access to food and water. All experimental procedures were approved by the Animal Experiments Ethics Committee of Fourth Military Medical University.

### 2.2. middle cerebral artery occlusion/ reperfusion

Middle cerebral artery occlusion/ reperfusion (MCAO/R) was induced using the intraluminal filament method [22]. Mice were anesthetized via 2% isoflurane inhalation (RWD Incorporated, Shenzhen, China). After the left common carotid artery (CCA) and external carotid artery were isolated and ligated, a silicone-coated monofilament (Stone persimmon, Zhengzhou, China) was inserted into the internal carotid artery through an incision in the CCA and reached the origin of the middle cerebral artery. The monofilament was withdrawn to restore blood flow 60 min after occlusion. Body temperature was maintained at 37°C throughout the procedure. Regional cerebral blood flow (CBF) was monitored before and after MCAO/R using a laser speckle imaging system (RFLSI III, RWD Incorporated).

### 2.3. Cell culture and treatment

Primary microglia and astrocytes were isolated from the brains of neonatal mice [23]. Brain cortices were dissected and placed in Ca^2+^- and Mg^2+^-free ice-cold Hanks’ Balanced Salt Solution (HBSS) (Thermo Scientific, Waltham, MA, USA). The meninges were carefully removed, and cortices were incubated in 0.125% deoxyribonuclease (Sigma-Aldrich) and 0.125% trypsin (Gibco, Waltham, MA, USA) for 20 min at 37°C. The tissue was digested, and the cell suspension was filtered through a 70 µm cell strainer (Corning, New York, NY, USA). The cells were centrifuged at 300 *g* for 5 min at 4°C, and the pellet was resuspended in Dulbecco’s modified Eagle’s medium (DMEM) F12 (Invitrogen, Carlsbad, CA, USA). They were plated into poly-D-lysine pre-coated flasks (Solarbio, Beijing, China). After the mixed glial cells reached confluence, the flasks were placed on an orbital shaker (BSI-101, Being, Shanghai, China) and spun at 80 *g* for 6 h at 37°C. The cells that still adhered to the flasks were astrocytes, while microglia were enriched in the supernatant and then cultured for 24 h to allow adhesion. Microglia were treated with 4-hydroxytamoxifen (0.02 mg/mL, Sigma-Aldrich) for 48 h to induce Cre-loxP recombination.

Primary neurons were isolated from the cerebral cortices of embryos as previously described [24]. Cortices were dissected, dissociated with 0.125% trypsin (Gibco). Resuspended cells were cultured in neurobasal medium (Thermo Scientific) supplemented with 2% B27 and 1% glutamine for 7 days, half of the medium was replaced every 3 days.

Isolation of microglia from adult brain tissue was performed by using a magnetic-activated cell sorting (MACS) kit (Biolegend, San Diego, CA, USA), as previously described [25]. Brain single-cell suspensions at a concentration of 1 × 10^7^ cells/mL were prepared from peri-infarct area of mice 72 h after MCAO/R. After treated with Fc receptor blocking reagent (Elabscience, Wuhan, China) for 15 min, the cells were incubated with biotin-conjugated recombinant purinergic receptor P2Y (P2RY12) antibody (10 μL/10^6^ cells, MedChemExpress, Shanghai, China) at 4°C for 15 min and then incubated with streptavidin-conjugated nanobeads for 15 min. Magnetic separation was conducted utilizing mass spectrometry columns, and P2RY12^+^ cells were obtained. After centrifugation at 500 g for 5 min, the cells were resuspended in HBSS.

To investigate whether microglia affect the secretion of CXCL10 from astrocytes in vitro, microglia and astrocytes were co-cultured in a 0.4-µm pore size transwell chamber (Corning, NY, USA) [26]. Microglia were seeded in the upper chamber of transwell plate, astrocytes were plated onto the bottom chamber.

For primary T cells culture. Mice were euthanized and the spleen was removed and dissociated in phosphate buffer saline containing 2% fetal bovine serum (FBS, Sigma-Aldrich). After aggregates and debris were filtered, T cells were purified by using using anti-Cluster of differentiation (CD)3-conjugated magnetic beads (Miltenyi Biotec., Bergisch Gladbach, Germany), and the cells were cultured in roswell park memorial institute (Gbico) containing 1% FBS and cells were activated by co-treatment of anti-CD3 (5 µg/ml, Gbico) and anti-CD28 (5 µg/ml, Gbico) for 72 h before use.

The BV2 and U87 cells were cultured in DMEM (Invitrogen) supplemented with 10% FBS and 1% penicillin/streptomycin at 37 °C.

### 2.4. Single-cell Fura-2 Ca^2+^ imaging

Single-cell Fura-2 Ca^2+^ was detected in primary cultured microglia subjected to normoxia or OGD/R condition as previously described [27]. Microglia were washed with HBSS (Gibco) containing 1.3 mM Ca^2+^, then loaded with 2.5 μM Fura-2 (MedChemExpress) for 30 min. The fluorescence signals, excited at 340 nm and 380 nm, were captured by a monochromator (Polychrome V, TILL-Photonics, Munich, Germany)-equipped inverted microscope with a Fluar objective. Intracellular Ca^2+^ concentration was detected by calculating the ratio of fluorescence intensity 340 and 380. Yoda1 (MedChemExpress) and Adenosine 5’-triphosphate (ATP, MedChemExpress) at a final concentration of 5 μM was added in the system to activate Piezo1 channel or induce Ca^2+^ release from endoplasmic reticulum.

### 2.5. Short interfering RNA (siRNA)-mediated knockdown of Pizeo1

To induce *piezo1* knockdown in BV2 cells, small interfering RNAs (siRNAs, Hippo Biotechnology, Hangzhou, China) were designed and synthesized (Table S1). The siRNAs were added in to the culture medium at a final concentration 160 nM with Lipofectamine 2000 (Invitrogen) and incubated for 48 h at 37 °C.

To overexpress Pizeo1, the full-length cDNA was cloned into pcDNA-3.1 vectors (GenePharma, Shanghai, China). Cells were seeded in culture dishes and grown until they reached 80% confluence. The selected plasmids were transfected using Lipofectamine 2000 (Invitrogen) and used for further research 24 h after transfection.

### 2.6. Oxygen-glucose deprivation and regeneration

Oxygen-glucose deprivation and regeneration (OGD/R) were performed as follows. The growth medium was replaced by pre-warmed serum and glucose free DMEM (Invitrogen), then the cells were transferred into a hypoxic humidified incubator (27310, Stem Cell, Vancouver, Canada) at 37 °C, the hypoxic duration was 1 h for primary microglia and 2 h for BV2 cells based on previous reports [28, 29]. These cells were then transferred to their previous medium under normoxic conditions for 24 h.

### 2.7. Flow cytometry

Mice were deeply anesthetized. The peri-infarct area was harvested and enzymatically digested using a Neural Tissue Dissociation Kit (130-093-231, Miltenyi Biotec) according to the manufacturer’s instructions. Single-cell suspension was obtained by passing the tissue through a 70 μm cell strainer (Corning). The suspension was then diluted at a density of 10^5^ cells/100 µL and stained with fluorophore conjugated antibodies (Table S2) and the appropriate isotype controls. Flow cytometry were performed using LSR II flow cytometer (BD Biosciences, Jersey, NJ, USA) and data were analyzed with FlowJo software (BD Bioscience). Three-dimensional cytometry data were mapped onto two dimensions based on the t-distributed Stochastic Neighbor Embedding (t-SNE) algorithm by using the viSNE11 software.

### 2.8. Reverse transcription polymerase chain reaction

Total RNA was extracted by TRIzol (Invitrogen) and purity was assessed by using a nanodrop spectrophotometer (Thermo Scientific). cDNA was then synthesized by using a cDNA kit (Thermo Scientific). Reverse transcription polymerase chain reaction (RT-PCR) was run on a BioRad thermocycler (ProFlex, Thermo Scientific). PCR primers for the *Pizeo1*, *Cxcl10*, *Tnf-α*, *Ifn-γ*, *Interleukin* (*Il*)*-1β*, *Il-4*, *Il-6*, *Il-10*, *Glyceraldehyde-3-phosphate dehydrogenase* (*Gapdh*) were designed and optimized using PrimerDesign^©^ (Table S3). Raw threshold cycle values were normalized to that of the *gapdh*.

### 2.9. Neurological function assessment

Neurological deficits, sensorimotor asymmetry, somatosensory deficits and motor coordination were assessed by modified neurological severity score (mNSS), corner-turning test, adhesive removal test and rotarod test respectively at 24 h, 72 h, and 120 h after MCAO/R as previous described. All mice were trained for 3 days before MCAO/R injury. All neurological function assessments were performed by an investigator blinded to the group identity.

### 2.10. Infarct volume assay

Mice were euthanized with an overdose of 30% chloral hydrate, and the brains were quickly removed, frozen and manually sliced into 2.0 mm thick coronal sections. The sections were then incubated with 2.0% 2,3,5-tripenyltetrazolium chloride (TTC; Sigma-Aldrich) at 37°C in the dark, followed by fixation in 4% paraformaldehyde overnight. The sections were placed on a black background board and photographed using a camera (EOS R, Canon, Tokyo, Japan) equipped with an F2.8 100 mm macro lens (Canon). A stable 5,500K color temperature light was applied using two flashlights (V1, Godox, Zhuhai, China).

### 2.11. Evans Blue staining

Evans Blue dye was used as a tracer to measure blood-brain barrier permeability 72 h after MCAO/R. A solution of 2% Evans Blue (2 ml/kg, Sigma-Aldrich) was injected intravenously 2 h before mice were euthanized. Brains were removed, frozen and sliced into coronal sections of 2.0 mm thick. The sections were fixed in 4% paraformaldehyde overnight and then photographed. The Evans Blue staining volume was analyzed using Photoshop software (Version CC 2019).

### 2.12. Immunofluorescence staining

Brains were sliced into frozen sections of 20 μm thick. The sections were incubated with the primary polyclonal antibody against ionized calcium-binding adapter molecule-1 (Iba-1, 1:500, ab178846, Abcam, Cambridge, Britain), glial fibrillary acidic protein (GFAP, 1:500, ab68428, Abcam), neuron-specific nuclear protein (NeuN, 1:500, ab177487, Abcam), CXCL10 (1:500, ab306587, Abcam), Pizeo1 (ab-2844888, 1:500, Affinity Biosciences, Beijing, China) and CD3 (1:500, ab16669, Abcam) respectively, overnight at 4 °C. The sections were then incubated with the corresponding secondary antibody for 60 min at 37 °C. Nuclei were stained with 4’,6-diamidino-2-phenylindole (DAPI, Sigma-Aldrich) for 5 min. Immunofluorescence images were acquired by a fluorescent microscope (IX-71, Nikon, Tokyo, Japan).

### 2.13. Double staining of neuron-specific nuclear protein and terminal-deoxynucleotidyl transferase mediated nick end labeling

After labeling neurons with NeuN (1:500, ab177487, Abcam), Brain sections were stained using a terminal-deoxynucleotidyl transferase mediated nick end labeling (TUNEL) reagent with an In Situ Cell Death Detection Kit (Beyotime, Beijing, China). The nuclei were stained with DAPI (Sigma-Aldrich). Images were captured using an fluorescence microscope (IX-71, Nikon).

### 2.14. RNA sequencing analysis

Microglia were isolated from peri-infarct area of control or pizeo1^CKO^ mice 3 days after MCAO/R by MACS method. Total RNA was extracted, and both the quantity and quality were assessed. Paired-end sequencing libraries were performed using a mRNA NEBNext Poly selection Library Prep Kit (Illumina, Los Angeles, CA, USA). cDNA sequence was then conducted using the Illumina NovaSeq 6000 sequencing platform with multiplex paired-end 150 base pairs reads. The quality of sequence was evaluated with Fast QC (Version 0.11.5). Gene alignment and read quantification were analyzed using Illumina’s automated Dragen RNA-Sequencing pipeline. Differentially expressed genes, which is defined as transcript abundance > 1.00-fold while false discovery rate < 0.05, were analyzed with DESeq2 (version 1.36) using the Wald test in DESeq2, corrected for multiple comparisons.

### 2.15. T cell migration assay

T cell migration was assessed by a 0.4-µm pore size transwell chamber (Corning). A total of 1 × 10^6^ T cells were seeded in the upper chamber of transwell plate, and the bottom chamber was filled with conditional medium collected from microglia-astrocyte co-culture system. The transwell plate was maintained at 37°C for 16 h and T cell migration was quantified by assessing the number of T cells in bottom chamber via Flow cytometry.

### 2.16. Multiplex chemokine immunoassay

CXCL and C-C motif chemokine ligand (CCL) level in peri-infarct area was detected using a LEGENDplex™ 13-plex proinflammatory chemokine panel (740451, BioLegend). Whole brain was freshly dissected 72 h after reperfusion and peri-infarct area was separated with cold PBS. After homogenized and centrifugation for 10 min at 3600 g, the supernatants were collected and diluted in assay diluent buffer. The samples were incubated with the APC-conjugated capture beads and PE-conjugated detection reagents, then analyzed on Aria III and quantified using the LEGENDplex™ software, version 8.0.

### 2.17. Western blot

Peri-infarct area tissue or cultured cells were harvested and lysed with radioimmunoprecipitation assay lysis buffer (Solarbio, Beijing, China) containing phenyl methane sulfonyl fluoride (Sigma-Aldrich). Samples were electrophoretically separated on SDS-PAGE gels and then transferred onto polyvinylidene fluoride membranes (Millipore, Los Angeles, CA, USA). After blocking with 5% nonfat milk (Maxigenes, Sydney, Australia), the membranes were then incubated with primary antibodies, included anti-TNF-α (1:500, ab183218, abcam), CXCL10 (1:500, ab306587, Abcam), Pizeo1 (ab-2844888, 1:500, Affinity Biosciences, Beijing, China), β-actin (ab8227, abcam) overnight at 4 °C. The blots were then incubated with the species-appropriate secondary antibodies (ab205719, ab6721, abcam) for 1 h at room temperature. Finally, the bands were viewed with densitometry (Bio-Rad, Heracles, CA, USA) and then quantified with the ImageJ software.

### 2.18. Statistical analysis

Statistical analysis was performed using GraphPad Prism 8.0 software. For comparisons between two groups, unpaired Student’s t-test was used for continuous variables with normal distribution, Mann-Whitney test was performed for non-normally distributed data. For comparisons between multiple groups, one-way or two-way analysis of variance (ANOVA) followed by appropriate post hoc tests was used for continuous variables with normal distribution, and the Kruskal-Wallis test followed by Dunn’s post hoc test was used for non-normally distributed data. Differences were considered statistically significant when P < 0.05.

## 3. Results

### 3.1. CIRI induces Piezo1 activation and upregulation in microglia

To investigate the role of Piezo1 in CIRI, we first isolated and cultured microglia, astrocytes, and neurons from wild-type mice and subjected them to OGD/R, RT-PCR results showed that *Piezo1* expression was dramatically elevated in microglia (Fig. 1A) but was only slightly increased in astrocytes (Fig. 1B) and neurons (Fig. 1C) upon OGD/R condition compared with normoxia condition. We then examined *Piezo1* expression in MACS microglia, *Piezo1* mRNA level was significantly higher in MACS microglia isolated from mice subjected to MCAO/R compared with sham operation (Fig. 1D). Western blot analysis also confirmed the substantial upregulation of Piezo1 expression in MACS microglia at 24 h and 72 h after MCAO/R (Fig. 1E, F). To examine whether Piezo1 might mediate Ca^2+^ influx in microglia after stroke, we performed the single-cell Ca^2+^ imaging of fura-2 in primarily cultured microglia. Microglia displayed significantly increased fura-2 signal upon OGD/R condition compared with normoxia condition (Fig. 1G, I) in response to Yoda1, a Piezo1 chemical activator [30]. This response was completely abolished by removing extracellular Ca^2+^, indicating that Piezo1 mediates Ca^2+^ influx rather than Ca^2+^ release from the endoplasmic reticulum store upon OGD/R condition (Fig. S1). Microglia under both normoxia and OGD/R conditions responded similarly to ATP, which induces Ca²⁺ release from the endoplasmic reticulum, suggesting that OGD/R injury did not impair ATP-induced Ca²⁺ signaling (Fig 1H, I). For immunohistochemical staining of brain sections, Piezo1 immunoreactivity was abundantly enhanced in peri-infarct microglia after MCAO/R (Fig. 1J). The proportion of Piezo1 positive microglia in the peri-infarct area was increased from 5.5% in sham-operated mice, to 26.7% and 82.7% in MCAO/R injured mice on 24 h and 72 h, respectively (Fig. 1K). The proportion of Piezo1 positive microglia was also slightly increased within the infarct core area 72 h after MCAO/R (Fig. S2). Together, these results indicated that CIRI elicited prominent Piezo1 expression elevation and activation in microglia, suggesting a potential role of Piezo1 signaling in the pathogenesis of acute ischemic stroke.

**Fig. 1.**
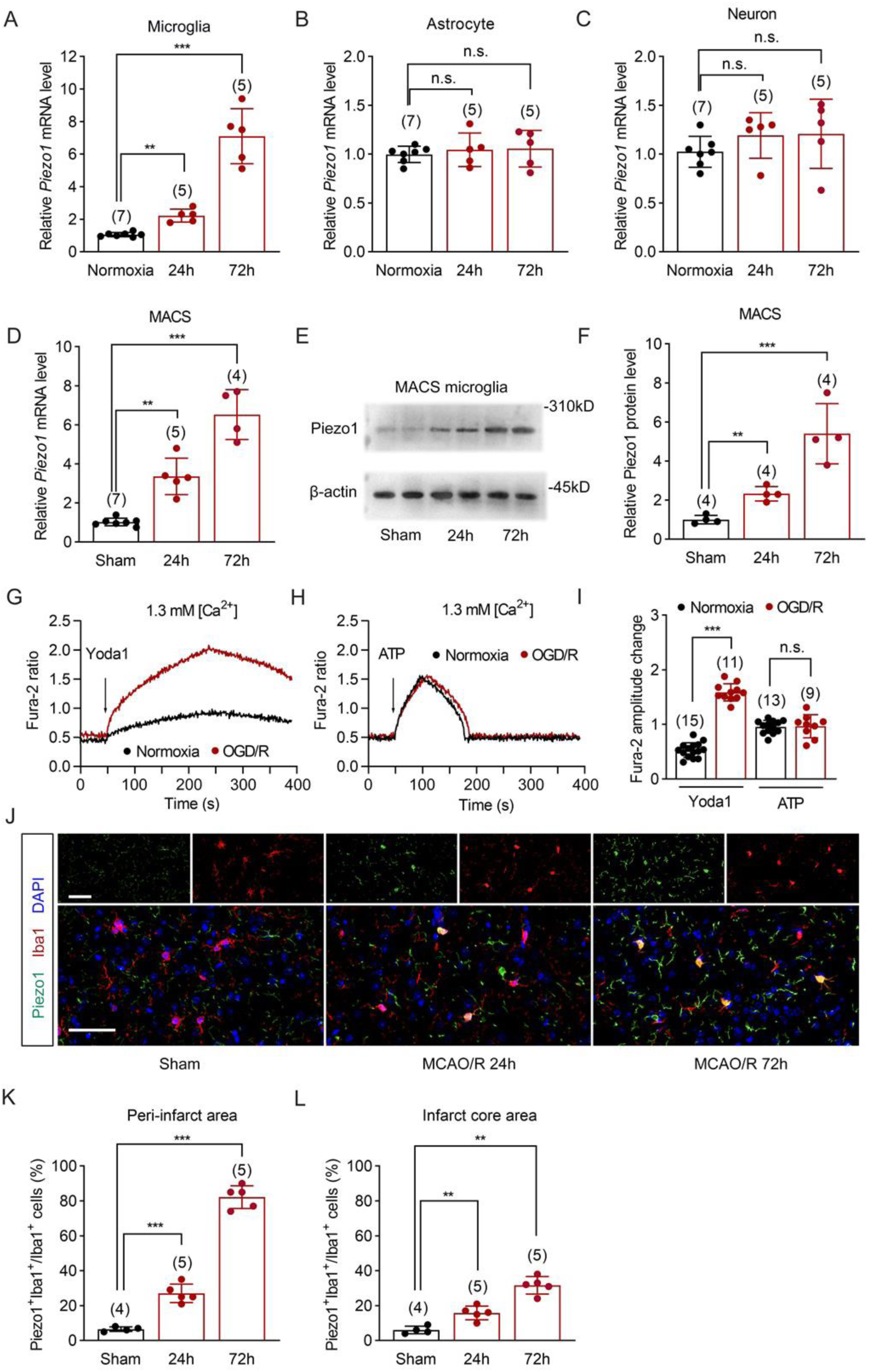
Piezo1 is upregulated and activated in microglia after CIRI. **A-C** RT-PCR analysis of relative *Piezo1* mRNA expression in primary microglia (A), astrocyte (B) and neuron (C) upon normoxia and OGD/R condition, with normoxia set to 1.0 (*n* = 4 per group). **D** RT-PCR analysis of the relative *Piezo1* mRNA expression in the MACS microglia isolated from mice subjected to sham operation or MCAO/R injury, with sham brain set to 1.0. **E, F** Western blot analysis of Piezo1 in MACS microglia isolated from mice subjected to sham operation or MCAO/R injury. A representative image was shown in E panel. Quantification of Piezo1 protein in indicated groups was shown in F panel by taking sham brain set to 1.0. **G, H** Representative fura-2 Ca^2+^ imaging of primary cultured microglia upon Normoxia or OGD/R condition response to Yoda1 (G) and ATP (H). **I** Scatterplot of maximal fura-2 amplitude changes in response to Yoda1 or ATP in indicated groups. **J** Representative immunofluorescence images of Piezo1 with Iba1 in the peri-infarcted regions after MCAO/R. Scale bar, 50 µm. **K, L** Quantification of the percentage of Piezo1^+^ cells among Iba1^+^ cells in peri-infarcted regions (K), and infarct cores (L). Data are expressed as mean ± SD. One-way ANOVA with Dunnett’s post hoc test for A, B, C, K and L. Two tailed unpaired Student’s t test for I. Kruskal–Wallis test with Dunn’s post hoc test for D, F. \*\**P* < 0.01, \*\*\**P* < 0.001, n.s.: no significant.

### 3.2. Conditional genetic deletion of Piezo1 in microglia ameliorated brain injury after MCAO/R

To determine whether Piezo1 signaling in microglia would have a specific impact on stroke outcome, the *Piezo1* conditional knockout mice were generated by crossing *Piezo1*^f/f^ mice with *Cx3cr1*^CreER^ mice (Fig. S3A-B). MCAO/R surgery was performed 14 days after tamoxifen administration (Fig. 2A). The efficacy of *Piezo1* conditional knockout in microglia was assessed 72 h after MCAO/R, as Piezo1 is barely detected under homeostatic condition. Immunostaining of Pizeo1 with Iba1 in brain slices confirmed the efficient conditional Piezo1 ablation (Fig. S3C, D). Piezo1^CKO^ mice showed no significant difference in body weight compared with control mice (Fig. S3E). The numbers of cerebral microglia, astrocytes, and neurons in Piezo1^CKO^ mice were comparable to those in control mice (Fig. 3F, G). No significant differences were observed between two groups in CBF at baseline, during ischemia as well as reperfusion (Fig. S3J, K). However, post-stroke Piezo1^CKO^ mice displayed significantly less severe sensorimotor deficits in a series of behavioral tests, including the mNSS (Fig. 2B), foot fault test (Fig. 2C), and rotarod test (Fig. 2D). TTC staining experiments showed that Piezo1^CKO^ mice exhibited significantly smaller infarct volume than control mice 72 h after MCAO/R (Fig. 2E, F). Poststroke Piezo1^CKO^ mice also showed a lower number of apoptotic neurons in peri-infarct area as revealed by TUNEL/NeuN staining (Fig. 2G, H). Additionally, we replicated the behavior tests and TTC staining experiments in adult female mice. Similar results were observed, we found a reduction of behavior deficits as well as infarct lesions in female Piezo1^CKO^ mice compared to controls (Fig. S4), Together, these findings indicated that Piezo1 signaling in microglia is essential for exacerbating functional deficits and brain injury following cerebral ischemia.

**Fig. 2.**
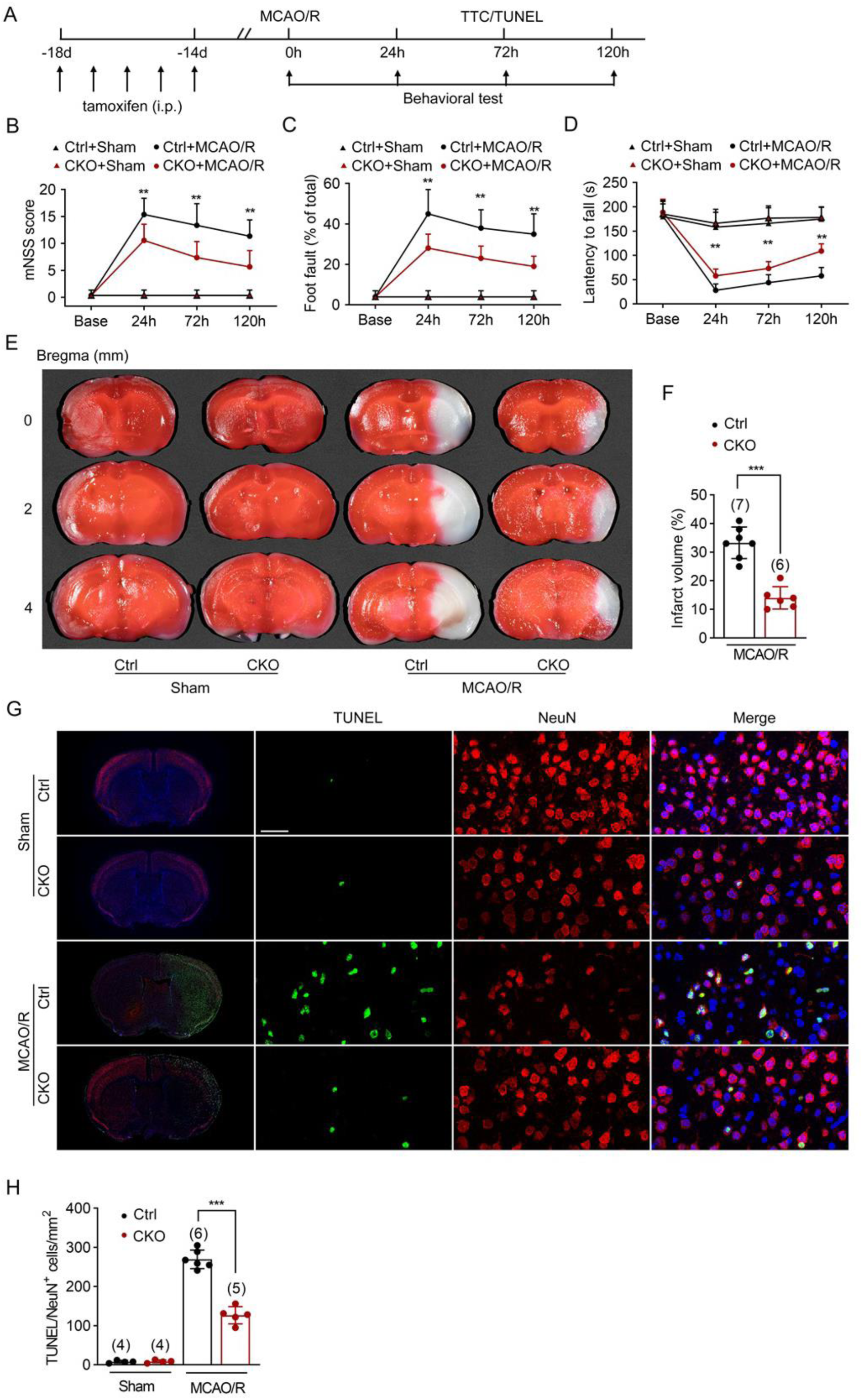
Piezo1 conditional knockout in microglia ameliorates poststroke outcomes. **A** Diagram of the experimental procedure. **B-D** Behavior deficits were assessed using the mNSS score (B), foot fault test (C), and rotarod test (D) in control and Piezo1^CKO^ mice 1, 3 and 5 days after MCAO/R. **E** Infarct volume was calculated by TTC staining in control and Piezo1^CKO^ mice. Representative three serial rostro-caudal sections from one mouse in indicated groups were displayed. **F** Quantitative analysis of the infarct volume in the indicated groups. **G** Apoptotic neurons were detected by TUNEL (Green)/NeuN (Red) immunostaining. Scale bar, 40 µm. **H** Quantitative analysis of the TUNEL/NeuN positive cells in indicated groups. Each symbol represents one mouse. Data are expressed as mean ± SD and analyzed by two-way repeated measures ANOVA with Greenhouse–Geisser correction followed by Sidak’s post hoc test for B, C and D. Two-tailed unpaired t-test for F. One-way ANOVA followed by Dunnett’s post hoc test for H. ** *P* < 0.01, *** *P* < 0.001.

**Fig. 3.**
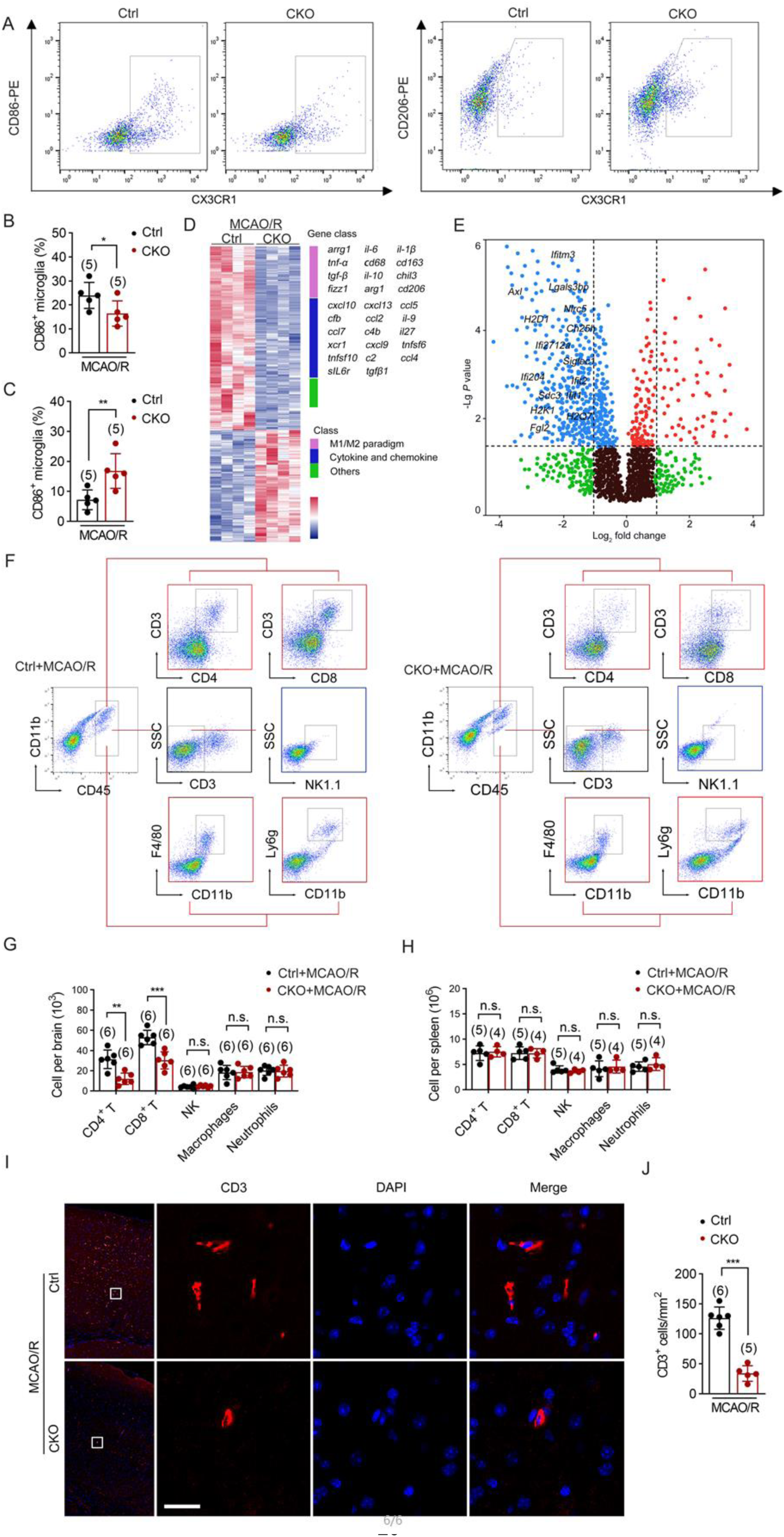
Piezo1 deficiency in microglia aggravates poststroke T cell CNS infiltration. **A** CD11b^+^CD45^int^ cells from the peri-infarct area of brain 72 h after MCAO/R were sorted using flow cytometry with anti-CX3CR1/CD86 and anti-CX3CR1/CD206. **B-C** Quantitative analysis of the percentage of CD86^+^ microglia (B) and CD206^+^ microglia (C). D Heatmap of differential gene expression profiles of MACS microglia from control and Piezo1^CKO^ mice 72 h after MCAO/R. E Volcano plot analysis displaying differentially expressed genes related to disease-associated microglia in MACS microglia caused by Piezo1 deficiency. Differentially expressed gene is defined as log_2_ (fold change) > 1.0 or < −1.0 meanwhile *P* vaule < 0.05. F Flow cytometry analysis of infiltrating lymphocytes subsets in brain of control and Piezo1^CKO^ mice 72 h after MCAO/R. G-H The infiltrating lymphocytes number in peri-infarct area (G) and spleen (H) in the indicated groups 72 h after MCAO/R. I Representative immunofluorescence images of CD3 in peri-infarct area of indicated groups after MCAO/R. Scale bar, 50 µm. J Quantitative analysis of the CD3 cells number in indicated groups. Each symbol represents one mouse. Data are expressed as mean ± SD. Two-tailed unpaired t-test for B, C, G, H and J. \**P* < 0.05, ** *P* < 0.01, *** *P* < 0.001, n.s.: no significant.

### 3.3. Piezo1 deficiency in microglia aggravates T cell infiltration

We assessed the impact of Piezo1 deficiency on microglia 72 h after MCAO/R. Flow cytometry analysis showed that Piezo1 deficiency in microglia decreased the proportion of CD86^+^ anti-inflammatory microglia (Fig. 3A, B) but increased the proportion of CD206^+^ anti-inflammatory microglia (Fig. 3A, C) in the peri-infarct area compared with control mice. We then performed RNA-sequencing on MACS microglia isolated from control and Piezo1^cKO^ mice 72 h after MCAO/R. *Piezo1* knockout resulted in differential expression of multiple genes related to microglial phenotypes, including *il-6*, *il-1β*, *tnf-α*, *cd68*, *arrg1*, *cd163*, *tgf-β*, *il-10*, *chil3*, *fizz1* (Fig. 3D). Notably, a number of microglial disease-associated signatures hallmark genes, including *axl*, *ank*, *apoe*, *cst7*, *cd9*, *cd63*, *ch25h*, *clec7a*, *ctsb*, *ctsd*, *ctsl*, *ctsz*, *fgl2*, *ifitm3*, *ifi27l2a*, *ifit1*, *ifit2*, *ifi204*, *itgax*, *lgals*, *lpl*, *siglec1*, *spp1*, *tyrobp*, *trem2* showed remarkable downregulation following Piezo1 ablation (Fig. 3E). These findings suggest that defective Piezo1 in microglia leads to its polarization towards an anti-inflammatory phenotype in response to cerebral ischemia. Previous studies have shown that activated microglia facilitates infiltration of peripheral leukocyte into brain, which further exacerbates brain inflammation and damage following stroke. Gene ontology analyses indicated that leukocyte infiltration was one of the most prominently altered biological processes (Fig. S5). Leukocyte accumulation in the peri-infarct areas of Control and Piezo1^CKO^ mice was measured after MCAO/R using flow cytometry. Deletion of Piezo1 in microglia considerably decreased brain infiltration of T lymphocytes (CD3^+^CD4^+^ and CD3^+^CD8^+^) (Fig. 3F, G), but did not impact the lymphocytes number in the spleen (Fig. 3H). Immunostaining of brain slices also revealed decreased T cell recruitment in Piezo1^cKO^ mice 72 h after MCAO/R compared with control mice (Fig. 3IJ). When T were depleted from the blood (Fig. S6A), T cell infiltration in peri-infarct areas (Fig. S6B), mNSS scores(Fig. S6C), foot fault rates (Fig. S6D), latency to fall time in rotarod test (Fig. S6E), and infarct volumes (Fig. S6F, G) were comparable between control mice and Piezo1^cKO^ mice after MCAO/R. Together, these results suggest that the loss of Piezo1 in microglia reduces T cell accumulation in the ischemic brain, exerting a protective effect against CIRI.

### 3.4. Piezo1 deficiency in microglia aggravates T cell infiltration via CXCL10 induction

Peripheral leukocyte recruitment substantially depends on locally elevated chemokines in the ischemic brain [5]. To investigate the role of Piezo1 in chemokine production, we measured chemokine levels in the peri-infarct area using a multiplex immunoassay. Piezo1 knockout in microglia significantly reduced the levels of CCL2, CCL8, CXCL9, and CXCL10 (Fig. 4A, B). We focused special attention on CXCL10, as it has been shown to be crucial for T cell infiltration by binding to CXC receptor 3, which is predominantly expressed on activated T cells [31]. Both mRNA and protein level of CXCL10 were further analyzed in peri-infarct area in control and Piezo1^CKO^ mice. In comparison to sham mice, higher *Cxcl10* mRNA level was detected as early as 12 h after MCAO/R and increased by over 100 folds at 24 h and 72 h after reperfusion, this increase is ameliorated by Piezo1 deficiency in microglia (Fig. 4C). Western blot assay also indicated CXCL10 level was roughly 10 times less abundant in Piezo1^CKO^ mice than that in controls (Fig. 4D, E). Several researches reported microglia as a source of CXCL10 during multiple diseases in central nerves system [32, 33]. In peri-infarct area of mice 72 h after MCAO/R, CXCL10 immunostaining was barely observed in microglia but was predominantly detected in astrocytes (Fig. S7). Expectedly, Piezo1 deficiency in microglia significantly decreased the number of CXCL10 expressing astrocytes in stroke mice (Fig. 4F, G). Increased peripheral leukocyte infiltration could also arise from brain blood barrier leakage. Indeed, we found that Piezo1 deletion in microglia markedly ameliorated brain blood barrier injury after stroke, as evidenced by Evans blue staining (Fig. 4H, I). Overall, these findings suggested that Piezo1 ablation in microglia resulted in significantly decreased chemokine secretion and improved brain blood barrier integrity post-stroke, ultimately reducing CNS infiltration of lymphocytes, particularly T cells.

**Fig. 4.**
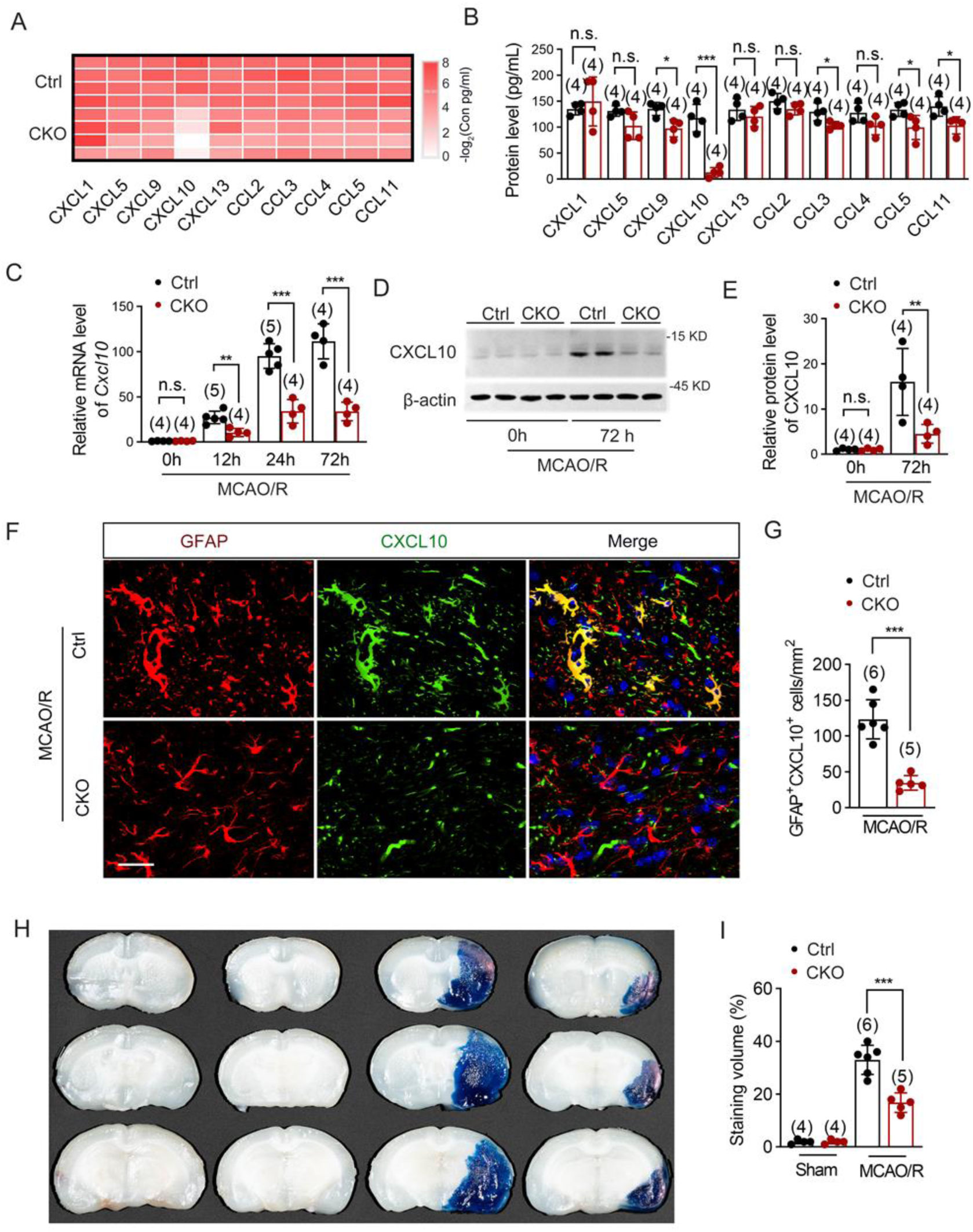
Piezo1 deficiency in microglia ameliorates T cell infiltration into ischemic brain via reducing CXCL10 induction. **A,B** Heatmap plot of multiplexed immunoassay showing chemokine expression profiles in the ischemic brains of indicated groups. Quantification of chemokine concentrations in indicated groups is shown in panel B, results were generated according to clustering of proteome profiler array assays of the listed chemokine. **C** *Cxcl10* mRNA levels detected by RT-PCR in peri-infarct area of Piezo1^CKO^ mice within the first 72 h after MCAO/R. **D.E** Western blot analysis of CXCL10 in peri-infarct area separated from control and Piezo1^CKO^ mice subjected to sham operation or MCAO/R injury. Reprehensive image was shown in panel D. Quantification of CXCL10 protein in indicated groups was shown in panel E, with sham brain set to 1.0. **F** Immunostaining for CXCL10 and GFAP of brain sections 72 h after MCAO/R. Scale bar, 50 µm. **G** Quantitative analysis of the GFAP and CXCL10 positive cells number in panel F. **H** Representative Evans blue staining images of brain sections. I Quantitative analysis of the Evans blue staining volume in panel H. Each symbol represents one mouse. Data are expressed as mean ± SD. Two-tailed unpaired t-test for B and G. Two-way ANOVA followed by Sidak’s post hoc test for C and E. One-way ANOVA followed by Dunnett’s post hoc test for I. \**P* < 0.05, ** *P* < 0.01, *** *P* < 0.001, n.s.: no significant.

### 3.5. Microglia modulate astrocyte CXCL10 production in a Piezo1 dependent manner

We next aimed to determine whether Piezo1 ablation in microglia is sufficient to block CXCL10 secretion from astrocytes. Primary cultured astrocytes were co-cultured with microglia isolated from brains of control or Piezo1^CKO^ neonatal mice (Fig. 5A). Upon OGD/R condition, we did detect lower level of CXCL10 in the co-culture media of Piezo1^CKO^ microglia than that of controls (Fig. 5B). Consistently, flow cytometry analysis showed that the CXCL10 ^+^ percentage was significantly less in astrocyte co-cultured with Piezo1^CKO^ microglia (Fig. 5C, D). Furthermore, we applied the co-culture conditional medium and T cells transmigration system to assess cell chemotaxis (Fig. 5E). A large number of migrated T cells were observed in response to the conditional medium from control group upon OGD/R condition, which is significantly decreased by Piezo1 ablation in microglia (Fig. 5F). We then applied antibody to block the action of CXCL10 in the transmigration system (Fig. 5G). Both ELISA and flow cytometry results showed that blocking CXCL10 reduced its levels (Fig. 5H) and equalized T cell migration (Fig. 5I) between Piezo1^CKO^ and control groups. This suggests that microglial Piezo1 induces T cell invasion by stimulating CXCL10 production in astrocytes following CIRI. Next, we assessed the impact of Piezo1 expression levels in microglia on astrocyte CXCL10 production. We co-cultured astrocyte with BV2 cells, in which Piezo1 expression could be manipulated (Fig. 5J, M). In accordance with primary cultured microglia, Piezo1 expression in BV2 cells was significantly increased following OGD/R injury (Fig. S8A, B). Transfection with Piezo1-specific siRNA (siPiezo1) reduced Piezo1 expression by 73% (Fig. S8C, D). Both CXCL10 concentrations, as well as the proportion of CXCL10-expressing astrocytes in co-cultures, were significantly decreased in the presence of Piezo1-silenced BV2 cells (Fig. 6K, L). On the other hand, *Piezo1* plasmid effectively stimulated an almost threefold increase in Piezo1 expression (Fig. S8E, F). CXCL10 concentrations (Fig. 6M) and the proportion of CXCL10-expressing astrocytes in co-cultures was increased in the presence of Piezo1-overexpressed BV2 cells (Fig. 6N, O). Together, these results indicated that following ischemia and reperfusion stimuli, microglial Piezo1 is a critical determinant in stimulating CXCL10 secretion from astrocyte.

**Fig. 5.**
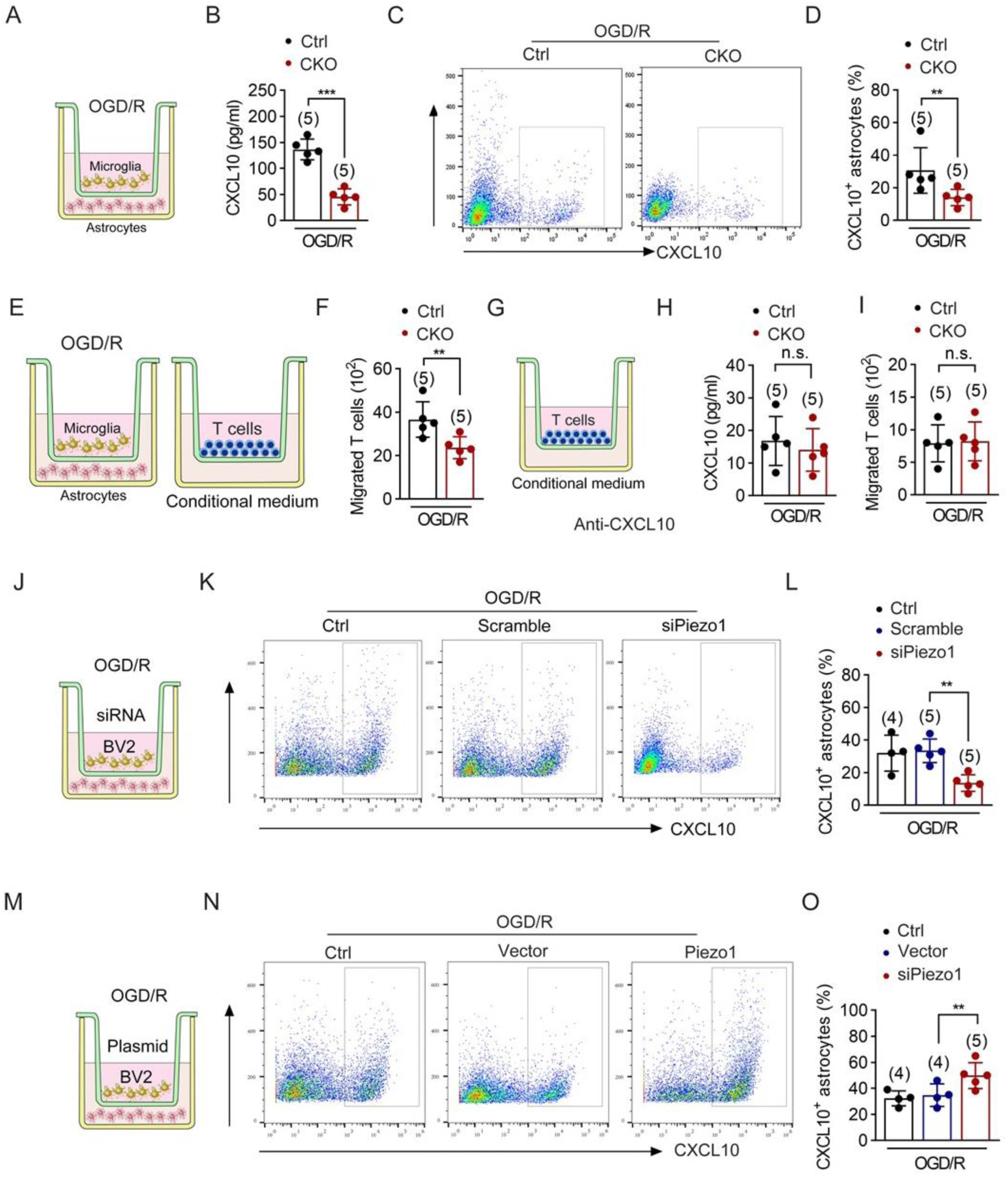
Microglia modulates astrocyte CXCL10 production in a Pizeo1-dependent manner. **A** Illustration of the *in vitro* astrocytes and microglia co-culture system. The system contains primary cultured astrocytes in the lower chamber and primary microglia isolated from control or Piezo1^CKO^ mice in the upper chamber. **B** CXCL10 concentrations detected by ELISA in the lower chamber of co-cultures. **C, D** CXCL10 positive astrocytes detected by flow cytometry in indicated groups. Representative plots gate is shown in panel C, quantitative analysis of CXCL10 positive astrocytes is shown in panel D. **E** Illustration of the *in vitro* transmigration system. The system contains conditioned media from co-culture system in the lower chamber and T cells in the upper chamber. **F** CD3 positive cells detected by flow cytometry in indicated groups. **G** Illustration of the transmigration system adding CXCL10 antibody. **H** Detection of CXCL10 level by ELISA in astrocytes of indicated groups treated by anti CXCL10 antibody. **I** CD3 positive cells detected by flow cytometry in indicated groups. **J** Illustration of the *in vitro* astrocytes and BV2 co-culture system. The system contains astrocytes and BV2 cells transfected with scramble siRNA or siPizeo1. **K, L** flow cytometry analysis of CXCL10 expression in astrocytes co-cultured with control or Piezo1 silenced BV2 cells. Representative plots gate is shown in panel K, quantitative analysis of CXCL10 positive cells is shown in panel L. **M** Illustration of the *in vitro* astrocytes and BV2 co-culture system, BV2 cells were transfected with empty vector or Piezo1 plasmid. **N, O** flow cytometry analysis of CXCL10 expression in astrocytes co-cultured with control or Piozo1 over-expressing BV2 cells. Representative plots gate is shown in panel N, quantitative analysis of CXCL10 positive cells is shown in panel O. Data are expressed as mean ± SD. Two-tailed unpaired t-test for B, D, F, H, and I. One-way ANOVA followed by Dunnett’s post hoc test for L and Q. ** *P* < 0.01, *** *P* < 0.001, n.s.: no significant.

**Fig. 6.**
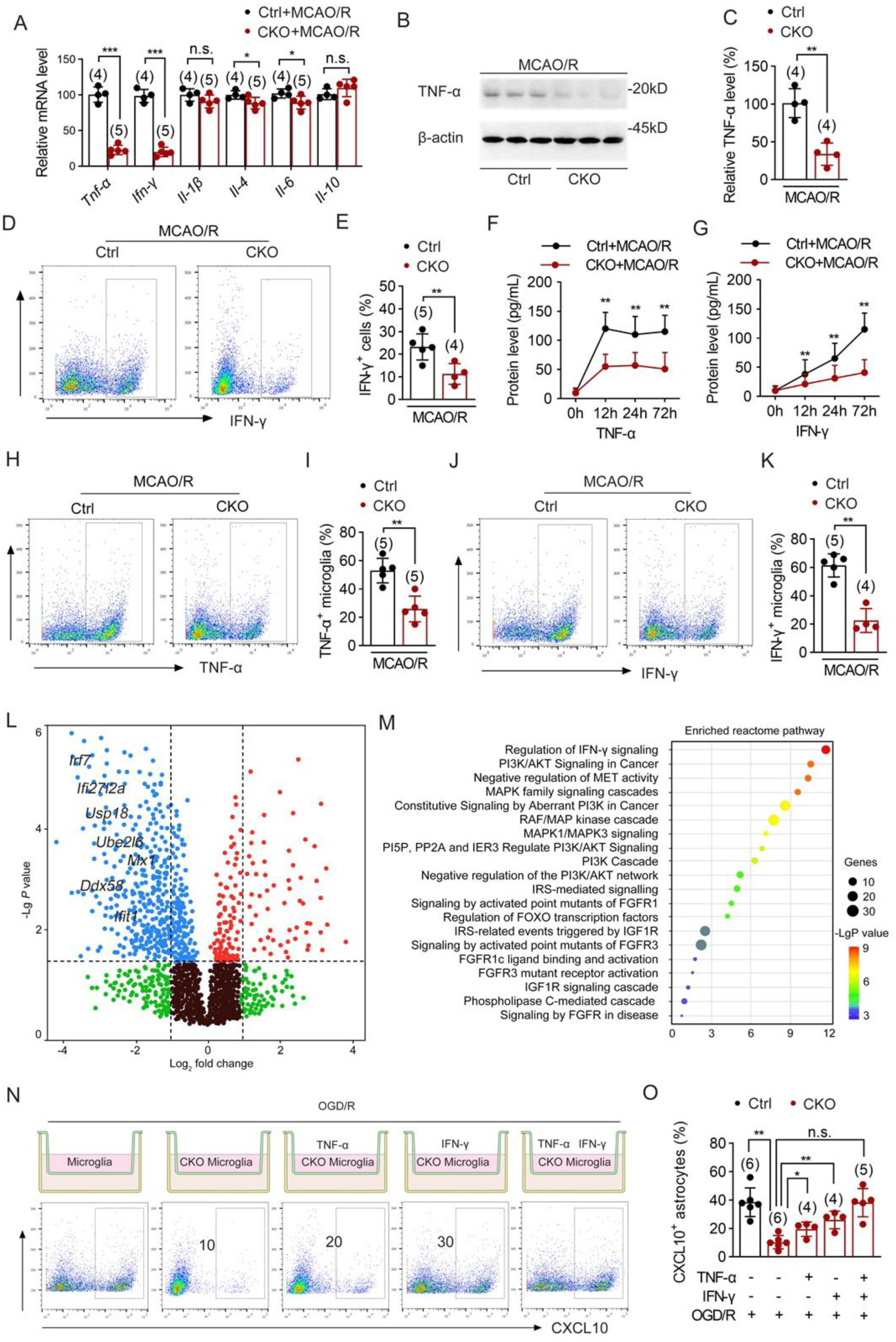
Microglia Piezo1 modulates astrocyte CXCL10 production via stimulating TNF-α and IFN-γ. **A** Relative cytokine mRNA levels in the peri-infarct area of control and Piezo1^CKO^ mice were detected by RT-PCR analysis 72 h after MCAO/R. **B** TNF-α in in peri-infarct area of control and Piezo1^CKO^ mice were detected by western blot analysis 72 h after MCAO/R. **C** Quantification of TNF-α of indicated groups, normalized to controls (set as 1.0). **D** flow cytometry analysis of IFN-γ positive cells in peri-infarct area of control and Piezo1^CKO^ mice 72 h after MCAO/R. **E** Quantitative analysis of IFN-γ positive cells in panel D. **F, G** *Tnf-α* (F) and *Ifn-γ* (G) mRNA level in peri-infarct area of control and Piezo1^CKO^ mice within the first 72 h after MCAO/R. **H** Flow cytometry analysis of TNF-α positive cells in MACS microglia isolated from peri-infarct area of control and Piezo1^CKO^ mice 72 h after MCAO/R. **I** Quantitative analysis of TNF-α positive cells in panel H. **H** Flow cytometry analysis of IFN-γ positive cells in MACS microglia isolated from peri-infarct area of control and Piezo1^CKO^ mice 72 h after MCAO/R. **I** Quantitative analysis of IFN-γ positive cells in panel H. **L** Volcano plot analysis showing dysregulated genes related to IFN-γ signaling performed in MACS microglia isolated from control and Piezo1^CKO^ mice at 72 h after MCAO/R. **M** Pathway enrichment analysis using Reactome datasets displaying the top 20 terms compared between control and Piezo1^CKO^ mice at 72 h after MCAO/R. **N** Flow cytometry analysis of CXCL10 positive astrocytes in astrocytes and control of Piezo1^CKO^ microglia co-culture system treated by TNF-α and/or IFN-γ antibodies upon OGD/R condition. **O** Quantitative analysis of CXCL10 positive cells in panel N. Data are expressed as mean ± SD. Two-tailed unpaired t-test for A, C, E, I and K. Two-way ANOVA with Sidak’s post hoc test in F, G. One-way ANOVA followed by Dunnett’s post hoc test for O. * *P* < 0.05, ** *P* < 0.01, *** *P* < 0.001, n.s.: no significant.

### 3.6. Microglia Piezo1 modulates astrocyte CXCL10 production via stimulating TNF-α and IFN-γ

Cytokine play synergistic roles in the induction of CXCL10 [34]. Accordingly, we measured the cytokine levels in the peri-infarct area of control and Piezo1^CKO^ mice 72 h after MCAO/R by RT-PCR. Notably, several key proinflammatory cytokines, including *Tnf-α*, *Ifn-γ*, were significantly decreased in Piezo1^CKO^ mice compared with controls (Fig. 6A). We paid interest in TNF-α and IFN-γ, because they are reported to play synergistic roles in the induction of CXCL10 [35]. Western blot analysis also indicated a decreased protein level of TNF-α in poststroke Piezo1^CKO^ brains compared with control mice (Fig. 6B, C). Furthermore, the percentage of IFN-γ positive cells was markedly less in poststroke Piezo1^CKO^ mice compared to controls (Fig. 6D, E). We next examined the dynamics of *Tnf-α* and *Ifn-γ* expression at earlier time points after MCAO/R. *Tnf-α* was upregulated and reached its maximum as early as 24 h post stroke, and maintained comparable levels at 24 h and 72 h in both groups (Fig. 6F). While upregulation of *Ifn-γ* displayed a time-dependent manner after MCAO/R in both groups (Fig. 6G). In sum, the absence of Piezo1 in microglia significantly ameliorated the inflammatory milieu in the acute phase of CIRI.

We then detected the impact of Piezo1 deletion on TNF-α and IFN-γ secretion in microglia following cerebral ischemia flow cytometry analysis gating on CD11b^+^CD45^int^ microglia. Piezo1^CKO^ mice exhibited less proportion of TNF-α (Fig. 6H, I) and IFN-γ (Fig. 6J, K) expressing microglia compared with control group. Accordingly, Piezo1 deficient BV2 cells also displayed significant lower level of *Tnf-α* and *Ifn-γ* gene level compared with control group following OGD/R injury (Fig. S9A). Notably, we observed a significant downregulation in the expression of IFN pathway-related genes including *Irf7*, *Ifi27l2a*, *Usp18*, *Ube2l6*, *Ddx58*, *Ifit1*, *Mx1* in Piezo1 deficient microglia (Fig. 6L). Further Reactome pathway enrichment analysis revealed Piezo1 knock out in microglia lead to prominent IFN-associated signaling changes (Fig. 6M). Together, these findings indicated critical roles of Piezo1 in secreting inflammatory cytokines in microglia after CIRI, particularly by activating IFN-γ signaling. In the co-cultures, administration of TNF-α or IFN-γ in medium of Piezo1 deficiency microglia increased CXCL10 positive astrocytes, while TNF-α and IFN-γ cotreatment resulted in a further increase in CXCL10 positive proportion (Fig. 6N). Together, these results suggested that TNF-α and IFN-γ mediated the crosstalk between microglia and astrocytes during CXCL10 induction, and this inflammatory response was remarkably intensified as a result of Piezo1 upregulation in microglia.

## 4. Discussion

Several studies have demonstrated an unequivocal association between Piezo1 dysfunction and worse outcomes of ischemic stroke in rodent model [21, 36, 37]. However, the underlying molecular mechanisms remains inadequately investigated. In this study, we provide novel insights into the mechanism by which piezo1 signaling robustly aggravates cerebral CIRI progression by modulating microglia initiated neuroinflammation. Overall, microglia Piezo1 is critical to aggravate secondary brain damage by increasing reperfusion induced TNF-α and IFN-γ, which could further increase production of chemokine CXCL10 by astrocytes and subsequent recruitment of T cells into the ischemic brain tissue.

Despite Piezo1 upregulation or dysfunction post stroke were observed in several reports, cell specific alteration of Pizeo1 expression and the resulting pathophysiological outcome remains elusive. In this study, we found that Piezo1 upregulation was more pronounced in microglia than in other cell types of brain after MCAO/R, suggesting that microglial Piezo1 may be a key driver of the excessive neuroinflammation seen in CIRI. Accordingly, both male and female microglia conditional Piezo1 deletion mice displayed ameliorated phenotypes with less behavior deficits and smaller infarct volume after MCAO/R. It should be noted that female mice displayed better outcome than those of genotype-matched males, probably owing to the beneficial effect of estrogen [38].

Transcriptome analysis revealed Piezo1 deficiency in microglia downregulated gene sets primarily involved in microglia phenotype. Gene oncology analysis indicated that microglial Piezo1 critically influences leukocyte infiltration in stroke pathophysiology. Similar alterations in microglia also contribute to neuroinflammation and disease progression in spinal cord injury [18, 39]. Infiltration of leukocyte in response to activation of microglia aggravates neuroinflammation and brain injury after CIRI [40]. Remarkably, we found defective microglia Piezo1 resulted in reduced infiltration of peripheral T cells, together with ameliorated disruption of the blood brain barrier. Whether these gene expression patterns regulated by microglia Piezo1 deficiency could impact poststroke long-term outcomes, such as cognitive decline, remains to be elaborated in further studies.

Further results showed that CXCL10, which is crucial for T cell infiltration, is increased in the peri-infarct area. Previous researches showed that microglia are the major source of CXCL10 [41]. In contrast to these findings, in our experiments, CXCL10 are primarily expressed in astrocytes in peri-infarct area. Possibly, astrocytes are crucial components of blood brain barrier [42]. CXCL10 from astrocytes could act in an autocrine or paracrine manner, drive cell injuries in itself and disruption of surrounding blood brain barrier components, that allows for subsequent T cells migration and recruitment. This mechanism gives a plausible explanation linking microglial Piezo1 activity, astrocytic CXCL10 production, and blood brain barrier breakdown.

Piezo1 deficiency in microglia decreased secretion of inflammatory cytokines in microglia and the whole brain, including TNF-α and IFN-γ. Microglia are major source of TNF-α in the early stage of CIRI in mice model [43, 44]. Accordingly, in our study, TNF-α reached its peak level as early as 12 h in the brain. While microglia are not the sole source of IFN-γ in the early phase of CIRI [45]. Typically, infiltrating T cells have been reported as an alternative source of IFN-γ at later stages of ischemic stroke [46, 47]. In line with these previous findings, we observed that IFN-γ level was continuous increased in the brain after stroke. Based on literature and our data, we hypothesize that microglia contribute to the initial surge in IFN-γ after MCAO/R, which then helps to recruit T cells; these T cells may in turn contribute further to IFN-γ production in the later phase. While our data implicate Piezo1 signaling in promoting TNF-α and IFN-γ production, likely through enhanced transcription, the exact molecular pathways linking Piezo1 activation to cytokine gene regulation remain to be determined.

## Ethics declaration

All animal experiments completely adhered to the ARRIVE guidelines and were approved by the Animal Care and Use Committee of Fourth Military Medical University.

## CRediT authorship contribution statement

Hui-Nan Zhang, Jing Huang, Tian Gao and Fei Liu wrote the main manuscript text. Hui-Nan Zhang, Hui-Min Chang, Bao Wang and Si-Jia Xu performed experiments and prepared figures. Fei Liu proof-read the manuscript. Hui-Feng Zhang and Xing Wang performed the data analysis. All authors reviewed the manuscript.

